# Disruption of Adaptive Immunity Enhances Disease in SARS-CoV-2 Infected Syrian Hamsters

**DOI:** 10.1101/2020.06.19.161612

**Authors:** Rebecca L. Brocato, Lucia M. Principe, Robert K. Kim, Xiankun Zeng, Janice A. Williams, Yanan Liu, Rong Li, Jeffrey M. Smith, Joseph W. Golden, Dave Gangemi, Sawsan Youssef, Zhongde Wang, Jacob Glanville, Jay W. Hooper

## Abstract

Animal models recapitulating human COVID-19 disease, especially with severe disease, are urgently needed to understand pathogenesis and evaluate candidate vaccines and therapeutics. Here, we develop novel severe disease animal models for COVID-19 involving disruption of adaptive immunity in Syrian hamsters. Cyclophosphamide (CyP) immunosuppressed or *RAG2* knockout (KO) hamsters were exposed to SARS-CoV-2 by the respiratory route. Both the CyP-treated and *RAG2* KO hamsters developed clinical signs of disease that were more severe than in immunocompetent hamsters, notably weight loss, viral loads, and fatality (*RAG2* KO only). Disease was prolonged in transiently immunosuppressed hamsters and uniformly lethal in *RAG2* KO hamsters. We evaluated the protective efficacy of a neutralizing monoclonal antibody and found that pretreatment, even in immunosuppressed animals, limited infection. Our results suggest that functional B and/or T cells are not only important for the clearance of SARS-CoV-2, but also play an early role in protection from acute disease.

**One Sentence Summary:** An antibody targeting the spike protein of SARS-CoV-2 limits infection in immunosuppressed Syrian hamster models.

## Main Text

The ongoing pandemic has led to the search for animal models faithfully recapitulating salient features of human coronavirus disease (COVID-19) for pathogenesis studies and evaluation of vaccines and therapeutics (*1*). Early reports indicated that Syrian hamsters, whose ACE2 is highly homologous to its human ortholog, were highly susceptible to SARS-CoV-2 infection but did not develop severe disease (*2, 3*). In 2008, administration of cyclophosphamide (CyP), an alkylating agent that suppresses B and T cell function, was used to develop a severe disease model for SARS-CoV in Syrian hamsters (*4*). We reasoned that a similar approach might allow the development of a severe disease model for SARS-CoV-2. Moreover, clinical findings of lymphopenia associated with COVID-19 (*5*) and its prominence in severe cases (*6, 7*) suggested an immunosuppressed animal model of COVID-19 might more accurately model some aspects of human disease. Here, we used CyP-treated and *RAG2* KO hamsters to investigate how transient disruption, or ablation, of the adaptive immune response affects SARS-CoV-2 infection. In addition, we used the immunocompetent and immunosuppressed hamster models to evaluate whether pre-exposure to virus through previous infection, or neutralizing antibodies, were sufficient to limit or prevent disease.

### Transient immunosuppression using CyP increased duration and severity of disease in SARS-CoV-2 infected hamsters

Wild-type hamsters are susceptible to SARS-CoV-2 infection and develop weight loss of approximately 10% 6 days post-infection (dpi) following a 100,000 plaque forming units (PFU) intranasal challenge dose (*2*). We confirmed this using both a 10,000 and 100,000 PFU dose (**Fig. S1A**). The maximum weight loss of each hamster was plotted for both a 10,000 and 100,000 PFU challenge dose demonstrating the high variability of wild-type hamsters in responses to virus infection with coefficients of variation of 57% and 48%, respectively (**Fig. S1B**). These hamsters developed a robust neutralizing antibody responses as measured by plaque reduction neutralization test (PRNT80) and virus genome was cleared 11 dpi, as measured by pharyngeal swab real time (RT)-PCR (**Fig. S1C,D**).

In our first experiment manipulating the adaptive immune response, we immunosuppressed hamsters using CyP (CyP-Tx) beginning −3 dpi. Suppression of lymphocytes was confirmed by hematology analysis just prior to challenge (**Fig. 1A**). On Day 0, groups of 10 hamsters were exposed to 100, 1,000, or 10,000 PFU of SARS-CoV-2 by the intranasal route. Disease progression was monitored by weight loss (**Fig. 1B**) and detection of viral RNA in pharyngeal swab using RT-PCR (**Fig. 1C**). Weight loss was remarkably similar between all SARS-CoV-2 exposed groups regardless of dose, with drastic weight loss beginning 6 dpi. Weight loss remained significant relative to starting weight through 35 dpi (**Table S1**). In contrast, mock (PBS) challenged hamsters treated with CyP steadily gained weight with a few exceptions (**Fig. S1C**). Specifically, one cage of four mock challenged hamsters showed evidence of weight loss 9-12 dpi (**Fig S2**). These same animals were the only mock challenged hamsters that were positive in the SARS-CoV-2 pharyngeal swab RT-PCR analysis starting 9 dpi (**Fig 1C** and **Fig S1C**).

**Fig. 1.**
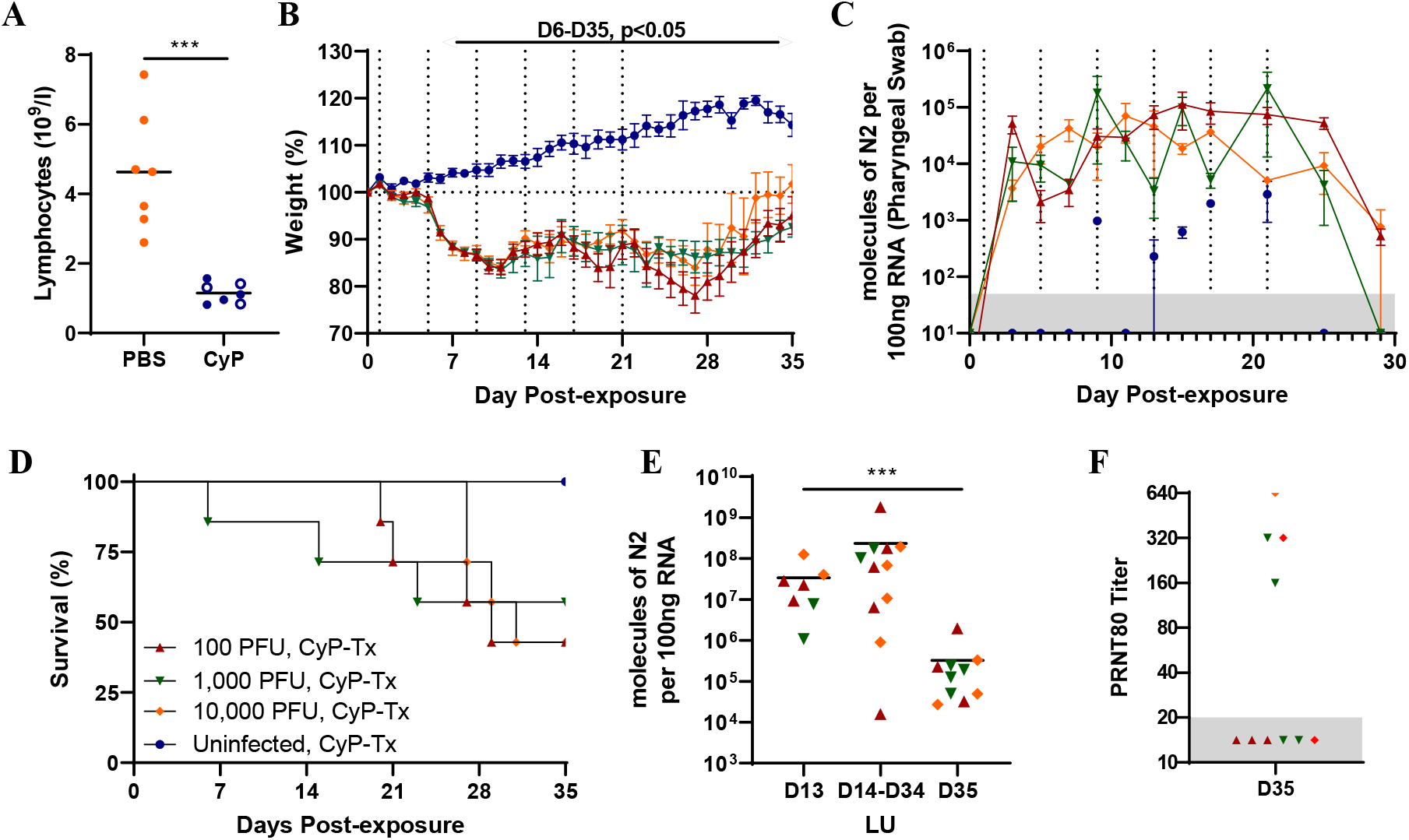
CyP-treated, SARS-CoV-2-infected hamsters. Groups of 10 Syrian hamsters each were immunosuppressed with CyP and **A)** Lymphocyte counts were determined from whole blood 3 days (closed symbols) or 4 days (open symbols) following CyP loading dose. CyP administration is depicted by vertical dotted lines in **B)** and **C)** and hamsters were exposed to increasing doses of SARS-CoV-2 intranally on Day 0. **B)** Weights were monitored for 35 days (statistical calculations in **Table S1**). **C)** Viral genome RNA copies per pharyngeal swab were assayed at indicated times post-infection. Hamsters were monitored for **D)** survival and **E)** lung tissue collected either at the time of death or scheduled euthanasia 13 or 35 dpi was assessed for viral load. **F)** Blood was collected from surviving hamsters 35 dpi and assessed by PRNT.

All of the SARS-CoV-2 challenged hamsters had detectable viral RNA in pharyngeal swabs at the first time point assayed, 3 dpi, and remained consistent (10^3^-10^5^ molecules of N2 per 100ng RNA) through the duration of CyP treatment (**Fig 1C**). After the last CyP treatment on 21 dpi, most of the hamsters continued to lose weight and several hamsters began to exhibit signs of a wasting disease, *i.e.* cachexia and extreme weight loss, requiring euthanasia (**Fig. 1B,D**). Then, approximately 7 days after cessation of CyP the remaining hamsters started gaining weight and viral RNA levels in pharyngeal swabs dropped below 10^3^ molecules of N per 100 ng RNA (**Fig. 1C**). Viral RNA was detected in lung tissue from a subset of hamsters collected13 dpi, the day of euthanasia of moribund animals (14-34 dpi), or euthanasia 35 dpi (end of study, **Fig. 1E**). There was a statistically significant reduction in lung viral burden comparing 35 dpi to 13 dpi lung homogenates. Serum collected from surviving hamsters 35 dpi was assayed for neutralizing antibodies (**Fig. 1F**). Six of the ten infected hamsters that survived in the CyP immunosuppression experiment never developed detectable levels of neutralizing antibodies. Interestingly, the four hamsters that did develop neutralizing antibodies after CyP treatment cessation rebounded more rapidly as measured by weight gain (**Fig. S3**). Together these data indicate that CyP treatment allowed a persistent infection that was reversed, in most animals, when CyP treatment was stopped and the adaptive immune response allowed to recover.

A second experiment was conducted to determine if a 3-day interval of CyP treatment might lead to even more dramatic weight loss than the 4-day interval. The last day of CyP treatment was 9 dpi rather than 21 dpi. Comparing the weights of ten hamsters treated with CyP at 3-day versus 4-day intervals revealed almost identical kinetics of weight loss (**Fig. S4A**). The cessation of CyP treatment9 dpi allowed more rapid recovery from weight loss. CyP treatment at 3-day intervals increased the amount of viral RNA detected by pharyngeal swab RT-PCR by approximately 1 order of magnitude relative to the 4-day interval between 3 and 7 dpi (**Fig. S4B**). The apparent increase level of virus did not result in increased weight loss or lethality. Based on these findings, we decided to use a 1,000 PFU intranasal dose and Cyp at a 4-day interval stopping 9 dpi for future antibody passive transfer experiments.

### *RAG2* KO hamsters infected with SARS-CoV-2 results in a lethal model

*RAG2* KO hamsters deficient in recombination activation gene 2 do not produce functional T or B cells (*8*). Whereas CyP treatment allowed transient and partial immunosuppression, the *RAG2* KO hamsters allowed us to evaluate the outcome of SARS-CoV-2 infection in the absence of functional lymphocytes. *RAG2* KO hamsters were exposed to 10,000 PFU SARS-CoV-2 by the intranasal route. The *RAG2* KO hamsters showed significant weight loss starting 3 dpi (**Fig. 2A**) and the infection was uniformly lethal with a median day-to-death of 6 days (p=0.0025, Log-rank test, **Fig. 2B**). Viral RNA detected from pharyngeal swabs trended to be higher than the CyP-treated hamsters, but not statistically significant (**Fig. 2C**). The 10,000 PFU and PBS mock groups from the CyP experiment shown in Fig. 1 were included in Fig. 2A-C for comparison. Organs from the *RAG2* KO animals that succumbed or met euthanasia criteria were homogenized and evaluated for viral RNA by RT-PCR. Viral RNA was present at the highest levels in the lung and trachea. Lower levels of viral RNA were also detected in extrapulmonary organs including the heart, liver, spleen, intestine, brain and kidney (**Fig. 2D**).

**Fig. 2.**
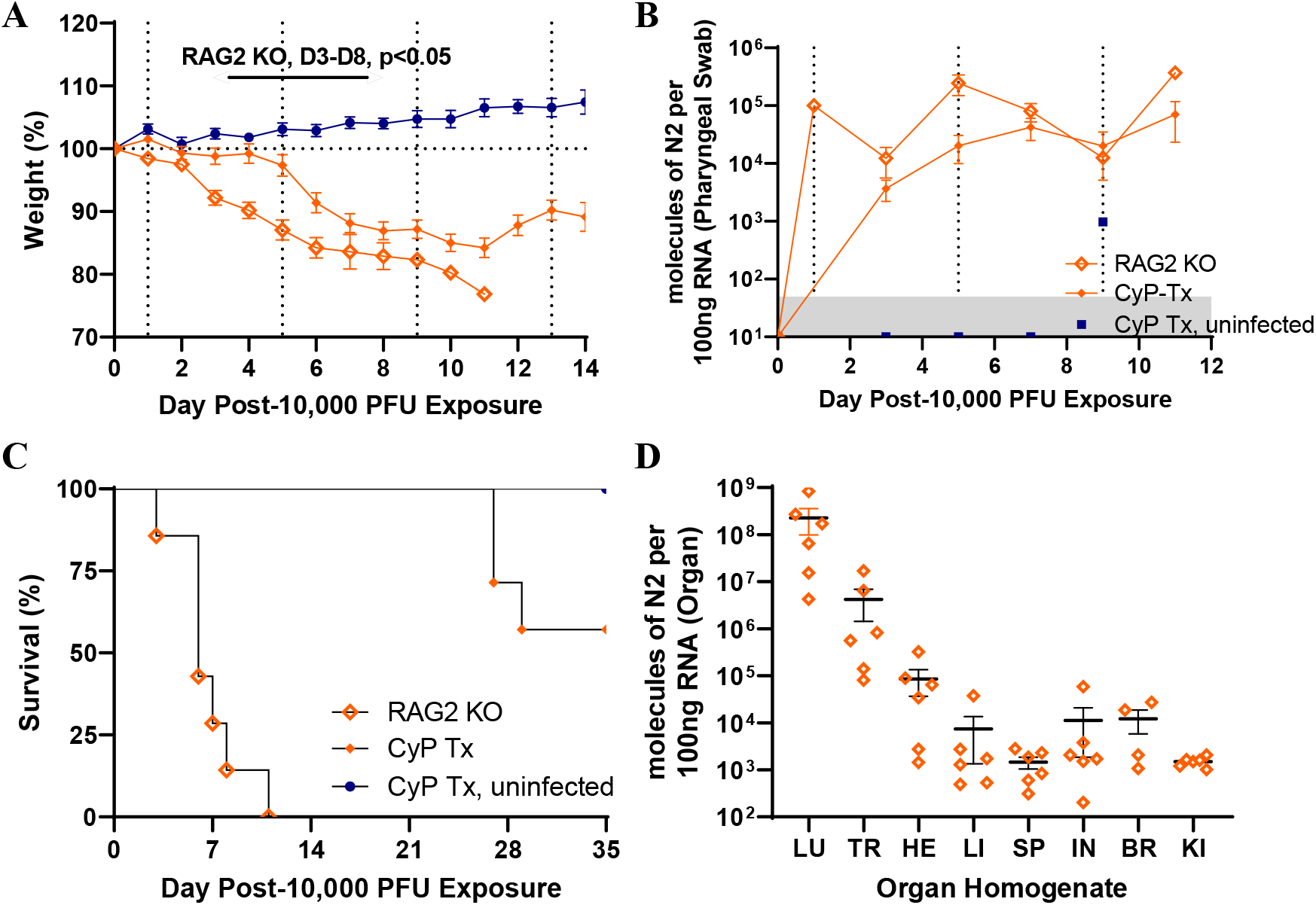
SARS-CoV-2-infected *RAG2* KO hamsters. Either *RAG2* KO (n=7) or CyP-treated hamsters (n=10) were exposed to 10,000 PFU SARS-CoV-2 on Day 0. Vertical dotted lines in **A)** and **B)** indicate CyP treatment for indicated animals. Hamsters were monitored for **A)** weight and **C)** survival. **B)** Viral RNA copies per pharyngeal swab were assayed at indicated times post-infection. **D)** Organs collected at the time of death were homogenized and assayed for viral load (LU=lung, TR=trachea, HE=heart, LI=liver, SP=spleen, IN=intestine, BR=brain, KI=kidney).

### Increased pathology of SARS-CoV-2-infected *RAG2* KO hamsters includes hemorrhage and severe edema

A subset of the SARS-CoV-2-infected hamsters from the CyP experiment shown in **Fig. 1** were euthanized 13 dpi. Lungs and nasal turbinates were formalin-fixed, paraffin embedded, and evaluated by histopathology and *in situ* hybridization (ISH) (**Fig. 3, Fig. S5**). Lung tissue from necropsied animals demonstrated focally extensive areas consolidation of pulmonary parenchyma admixed with dense aggregates of inflammatory cells (**Fig. 3A**). The bronchial lumina are multi-focally lined by hyperplastic respiratory epithelium (**Fig. 3B**) and SARS-CoV-2 genomic RNA was frequently detected in alveolar pneumocytes, alveolar infiltrates, and bronchiolar epithelial cells by ISH (**Fig. 3C**). At 14 dpi, immunocompetent hamsters infected with SARS-CoV-2 exhibited only mild congestion and inflammatory infiltration indicating recovery from viral challenge (*2*) supporting the hypothesis that CyP administration to disrupt adaptive immunity exacerbates and prolongs disease in hamsters.

**Fig. 3.**
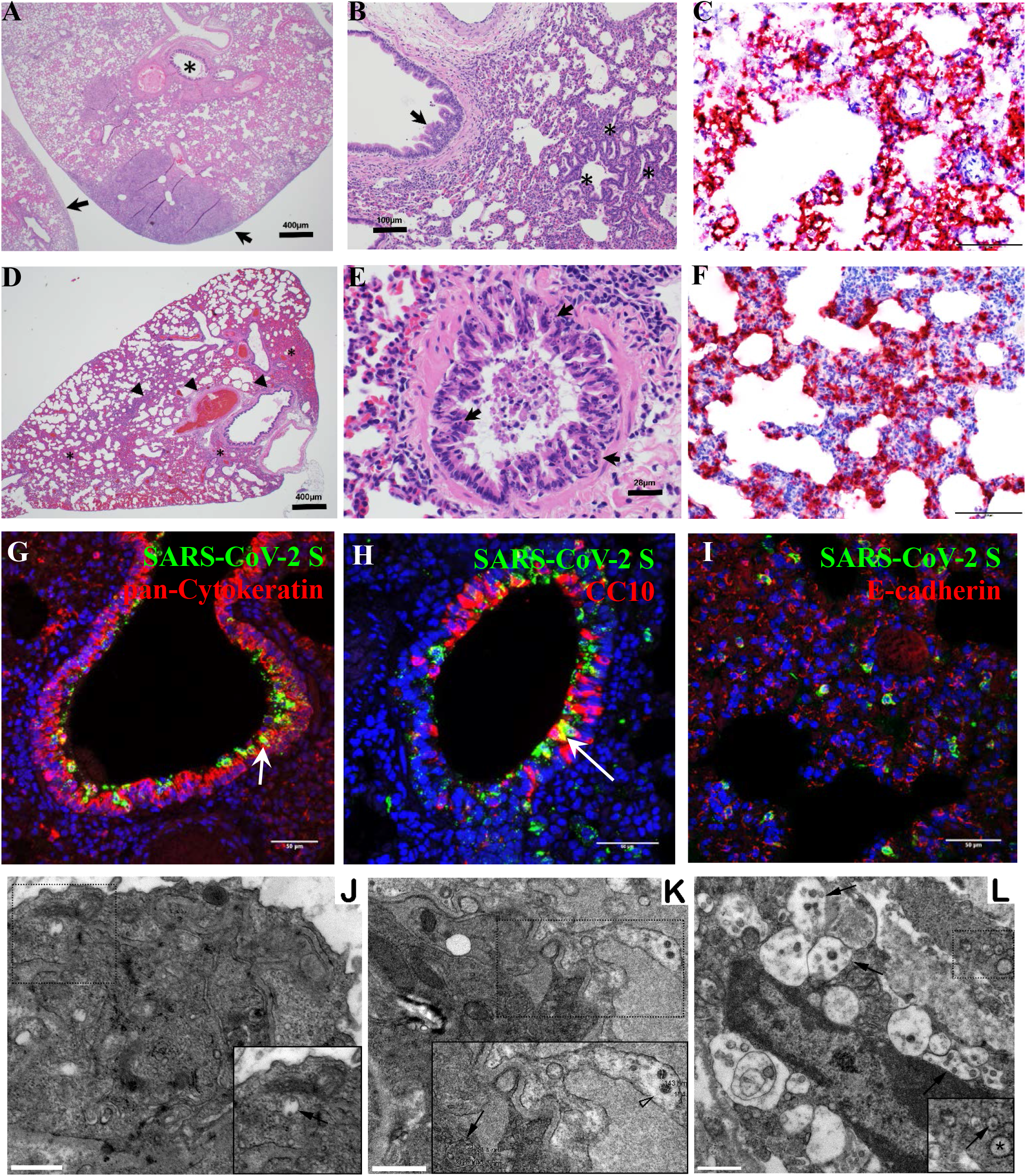
Pathology of SARS-CoV-2 in CyP-treated and *RAG2* KO hamsters. H&E sections (**A, B**) of lung tissue from CyP-treated hamsters euthanized 13 dpi show extensive areas of consolidation with dense aggregates of inflammatory cells. **A)** Bronchial lumen are lined by hyperplastic folds of respiratory epithelium (asterisk) and the pleural surface is multifocal thickened and expanded by fibrous connective tissue and inflammatory cells (arrows). **B)** The bronchial lumen are lined by hyperplastic folds of respiratory epithelium (arrow). Areas of alveolar septa lined rows of type 2 pneumocytes (asterisks). **C)** SARS-CoV-2 genomic RNA was frequently detected in alveolar pneumocytes, alveolar infiltrates, and bronchiolar respiratory epithelial cells from CyP-treated hamsters by ISH. H&E sections (**D, E**) of lung tissue from *RAG2* KO hamsters collected at the time of death. **D)** Areas of hemorrhage (asterisk) and inflammation (arrowheads) expanding the interstitium and connective tissue surrounding bronchi and arteries (arrows). **E)** Necrotic bronchial epithelium (arrows) overlaid by hemorrhagic exudate. Peribronchial connective tissue is expanded by lymphocytes, heterophils (asterisks) and fewer macrophages that often contain hemosiderin (arrowheads). There is marked consolidation in surrounding alveoli with marked septal congestion and expansion by previously mentioned inflammatory cells. **F)** SARS-CoV-2 genomic RNA was frequently detected in alveolar pneumocytes, alveolar infiltrates, and bronchiolar epithelial cells from *RAG2* KO hamsters by ISH. (**G-I**) Immunofluorescence assays demonstrate SARS-CoV-2 antigens (S or NP, green) were detected in brochiolar epithelium labelled anti-pan-cytokeratin antibody (red, **G**) club (clara) cells labelled by anti-CC10 antibody (red, **H**) and alveolar epithelial cells labelled by anti-E-cadherin antibody (red, **I**) in *RAG2* KO hamsters. (**J-L**) TEM of hamster lungs with increasing viral loads. **J**) Lung section from hamster with 10^6^ molecules of N2 per 100ng RNA. Inset shows cytoplasmic vacuole with possible virus (black arrow). **K**) Lung section from hamster with 10^7^ molecules of N2 per 100ng RNA. Potential mature viral particles (approximately 143-154nm diameter, arrowhead) are present at the cell periphery and suspected immature virions are detected more internally in a cytoplasmic vacuole (approximately 62-97nm diameter, black arrow). **L**) Lung section from hamster with 10^8^ molecules of N2 per 100ng RNA. Numerous cytoplasmic vacuoles of possible virus are evident (black arrows). Inset shows an example of swollen rER (asterisk) with virus forming within the swollen rER (black arrow). Scale bars (**A, D**) = 400 microns; (**B**) = 100 microns; (**E**) = 28 microns; (**C, F**) = 100 microns, (**G, H, I**) = 50 microns. (**J-L**) = 1 micron.

Lung tissue collected at the time of death of SARS-CoV-2-infected *RAG2* KO hamsters contain areas of hemorrhage and inflammation expanding the interstitium and connective tissue surrounding bronchi and arteries (**Fig. 3D**). As seen in the CyP-treated hamsters, the bronchial lumina of *RAG2* KO hamsters is lined with multiple layers of infected epithelial and club cells, containing infiltrates of lymphocytes (presumably nonfunctional), heterophils and macrophages (**Fig. 3E,G,H**). There is hemorrhage within alveolar lumen and consolidation with septal congestion. Also as observed in CyP-treated hamsters, SARS-CoV-2 genomic RNA and viral antigen were frequently detected in alveolar pneumocytes (**Fig. 3I**), alveolar infiltrates, and bronchiolar epithelial cells (**Fig. 3F**). Together these data indicate that the absence of functional B and T cells in *RAG2* KO hamsters allows increased pathology and lethality from SARS-CoV-2 infection.

Electron microscopy studies were performed on lung sections of SARS-CoV-2 infected, CyP-treated hamsters with varying lung viral loads (**Fig. 1**). The lungs lack a typical morphology wherein distinct alveolar, inter-alveolar septum with capillaries, Type I and Type II pneumocytes are clearly evident. The animal with the least viral load (10^6^ molecules of N2 per 100ng RNA, **Fig. 3J**) show some remnants of normal architecture while the animal with the highest viral load (10^8^ molecules of N2 per 100ng RNA, **Fig. 3L**) lack the prominent alveolar space and were congested with immune cells. The lamellar bodies in the Type II pneumocytes are still identifiable in hamsters with high viral loads.

SARS-CoV-2 is reported to range in diameter from 50-160nm (with surface spikes measuring approximately 20nm) (*9–13*). The low and high viral load samples show compartmentalized vacuoles with round virus-like structure and vesicles with irregular shaped structures, resembling multi-vesicular bodies (**Fig. 3J-L**). The more electron dense vesicles are suggestive of mature virus particles whereas, the less electron dense particles are likely immature virions. In addition to cytoplasmic vacuoles, these tissue show swollen rough endoplasmic reticulum (rER, **Fig. 3L**). Swollen rER appears as a result of cellular stress and/or increased viral protein synthesis. Immuno-gold labeling is necessary to confirm the presence of virus particles.

Tracheas from the same SARS-CoV-2 infected, CyP-treated hamsters with varying lung viral loads were analyzed by electron microscopy. These tracheas also exhibit disruption of the epithelial layer. Trachea from the animal with the lowest lung viral load (10^6^ molecules of N2 per 100ng RNA) showed mostly intact ciliated cells on the surface (**Fig. 4A**). These cells show several cytoplasmic vacuoles with potential immature virus particles (**Fig. 4D,E**). As lung viral load increases, the presence of ciliated cells and epithelial cells lining the tracheal lumen decrease (**Fig. 4B-C**, respectively). Very few ciliated cells were seen in the animals with the heaviest viral load (10^8^ molecules of N2 per 100ng RNA) and there is an absence of cytoplasmic content marked by the lack of electron dense cytoplasmic vesicles (**Fig. 4F**).

**Fig. 4.**
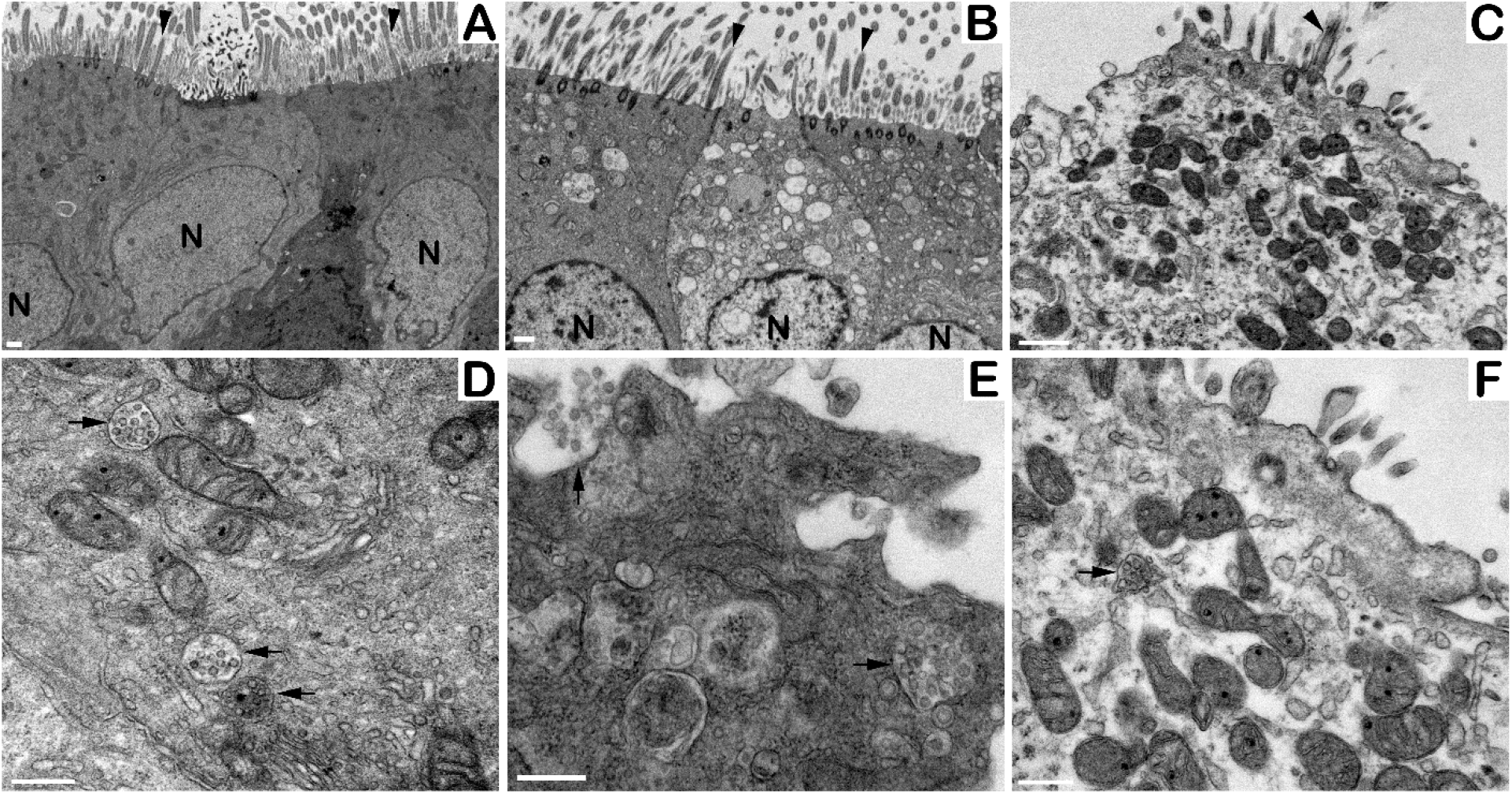
SARS-CoV-2 disrupts the tracheal epithelial layer. Tracheal sections were collected from SARS-CoV-2 infected, CyP-treated hamsters, sorted by lung viral load and analyzed by transmission electron microscopy. **A)** The animal with lung viral load of 10^6^ molecules of N2 per 100ng of RNA showed the most intact ciliated cells on the surface of the trachea (arrowheads). As viral load increases (from animals with lung viral load of 10^7^ (**B)** and 10^8^ (**C)** molecules of N2 per 100ng RNA, respectively) the presence of ciliated cells and epithelial cells lining the trachea lumen decrease. **D)** Cells from the low viral load animal show several cytoplasmic vacuoles with potential immature viral particles (arrows). **E)** The release of cytoplasmic vacuole content (possible immature virions, arrows) into the luminal space of a cell that has detached from the epithelial layer. **F)** From the highest viral load animal, very few ciliated cells are noted. Cytoplasmic vacuole with potential immature viral particles are observed (arrow). Scale bars (**A-C**) = 1 micron; (**D-F**) = 500 nm.

### Protective efficacy of prior SARS-CoV-2 infection

Nine previously infected immunocompetent hamsters from two separate experiments were retained for a re-challenge with SARS-CoV-2. Our goal was to determine if prior infection elicited protective immunity. To ensure that these hamsters had cleared infectious virus after the initial challenge, hamsters were treated with 3 doses of CyP and weights monitored for 14 days (**Fig. 5A**). If the animals were persistently infected, we predicted that hamsters would experience onset of disease as measured by rapid weight loss and positive pharyngeal swab RT-PCR following CyP administration. No significant weight loss was detected and swabs were negative for viral RNA (**Fig. 5A**). All nine animals were positive for neutralizing antibodies prior to the re-challenge (**Fig. 5B**). The nine previously exposed hamsters and seven naïve hamsters were challenged with 100,000 PFU SARS-CoV-2. Weights were monitored daily and pharyngeal swabs were collected every other day for viral RNA RT-PCR. Control animals lost weight as predicted whereas the previously exposed animals were protected from significant weight loss. Pharyngeal swab RT-PCR indicated a statistically significant reduction in viral RNA detected in re-challenged animals compared to naïve animals (**Fig. 5C, Table S3**). In addition, viral load in lung homogenates from animals euthanized at the end of the study indicated a statistically significant (p=0.0164, unpaired t test) reduction in viral RNA level in the re-challenged animals **Fig. 5D**). These data indicate that immunocompetent hamsters recovered from prior exposure to the SARS-CoV-2 does not result in persistent infection, but instead elicits protective immunity.

**Fig. 5.**
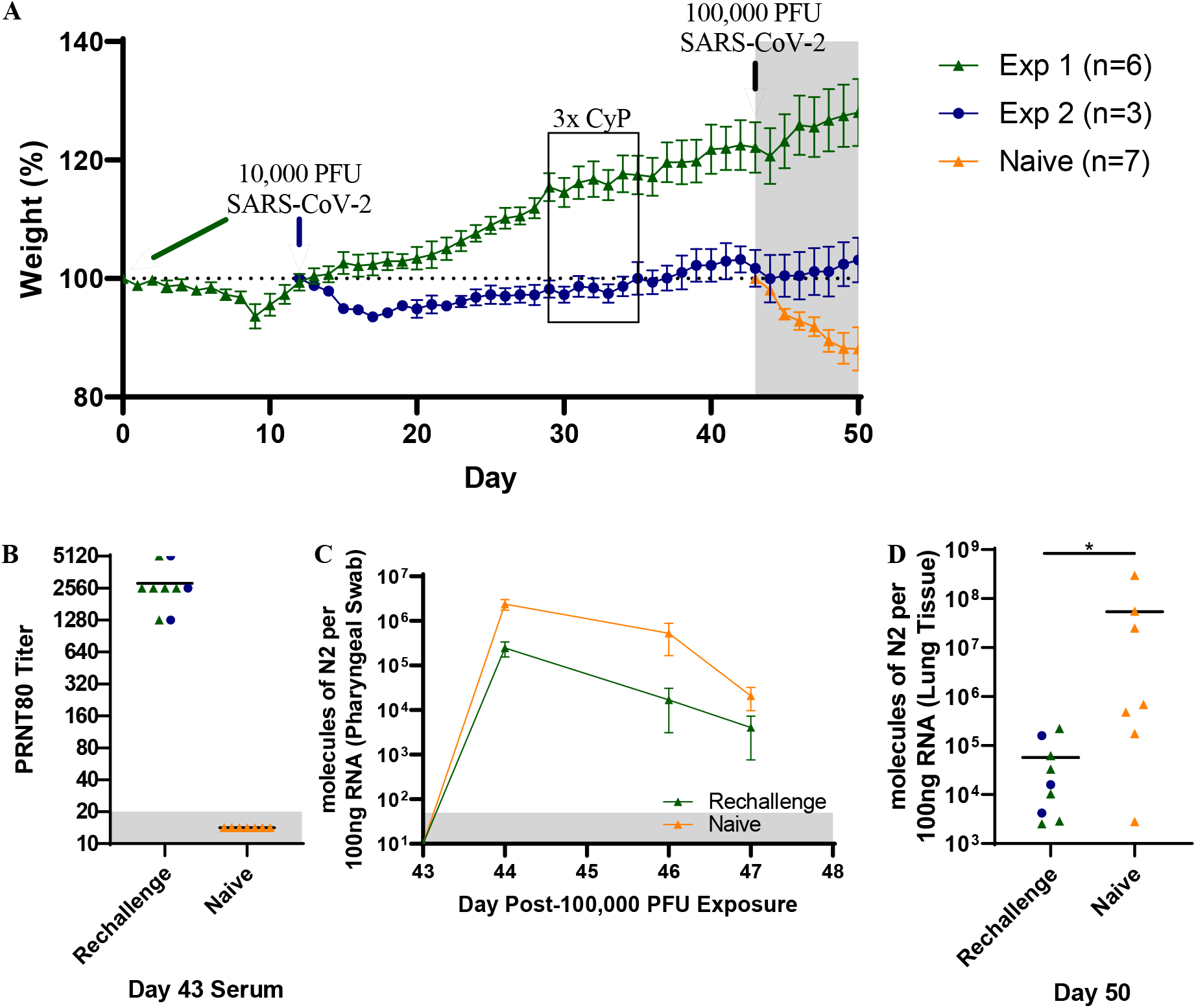
Re-challenge of previously infected SARS-CoV-2 hamsters. **A)** Weight data from hamsters initially exposed to 10,000 PFU SARS-CoV-2 and re-challenged with 100,000 PFU SARS-CoV-2. **B)** PRNT80 titers depicting the level of circulating neutralizing antibody prior to Day 43 virus exposure. Disease progression was monitored by **A)** weight and **C)** pharyngeal swabs. **D)** Lung tissue collected on Day 50 was assayed for viral genome.

### Protective efficacy of anti-SARS-CoV-2 mAb

A human monoclonal antibody (mAb) targeting the spike protein of SARS-CoV-2 was evaluated in a passive transfer experiment using immunosuppressed hamster. Thirty mg/kg of Centi-F1 mAb (IC50= 391 ng/mL, PRNT80=1280), a control IgG (normal) mAb, or PBS were administered −1 dpi. On Day 0 the immunosuppressed animals were challenged with 1,000 PFU SARS-CoV-2. Neutralizing antibody levels in sera at the time of challenge were measured by PRNT80 (**Fig. 6A**). Detectable levels ranging from 80 to ≥640 were found in the Centi-F1 mAb group. Disease progression was monitored by weight change (**Fig. 6B**) and pharyngeal swab RT-PCR (**Fig. 6C**). Hamsters administered the normal mAb or PBS exhibited significant weight loss starting 5 dpi and ultimately had group weights drop to 86% and 91%, respectively (**Table S4**). Hamsters Treated with Centi-F1 mAb maintained over 100% of their Day 0 weight to the conclusion of the experiment (13 dpi, **Table S4**). Virus RNA was detected in pharyngeal swabs in all groups but trending lower in Centi-F1 mAb-treated animals on Day 11 (p=0.1038, unpaired t test, **Table S4**). Similarly, lung tissue collected 13 dpi (end of study) indicate comparable levels of viral RNA detected (**Fig. 6D**), but significantly reduced infectious virus in Centi-F1 mAb-treated animals (p=0.0002, unpaired t test, **Fig. 6E)**. These findings demonstrate that a SARS-CoV-2 mAb can limit infection in immunosuppressed animals.

**Fig. 6.**
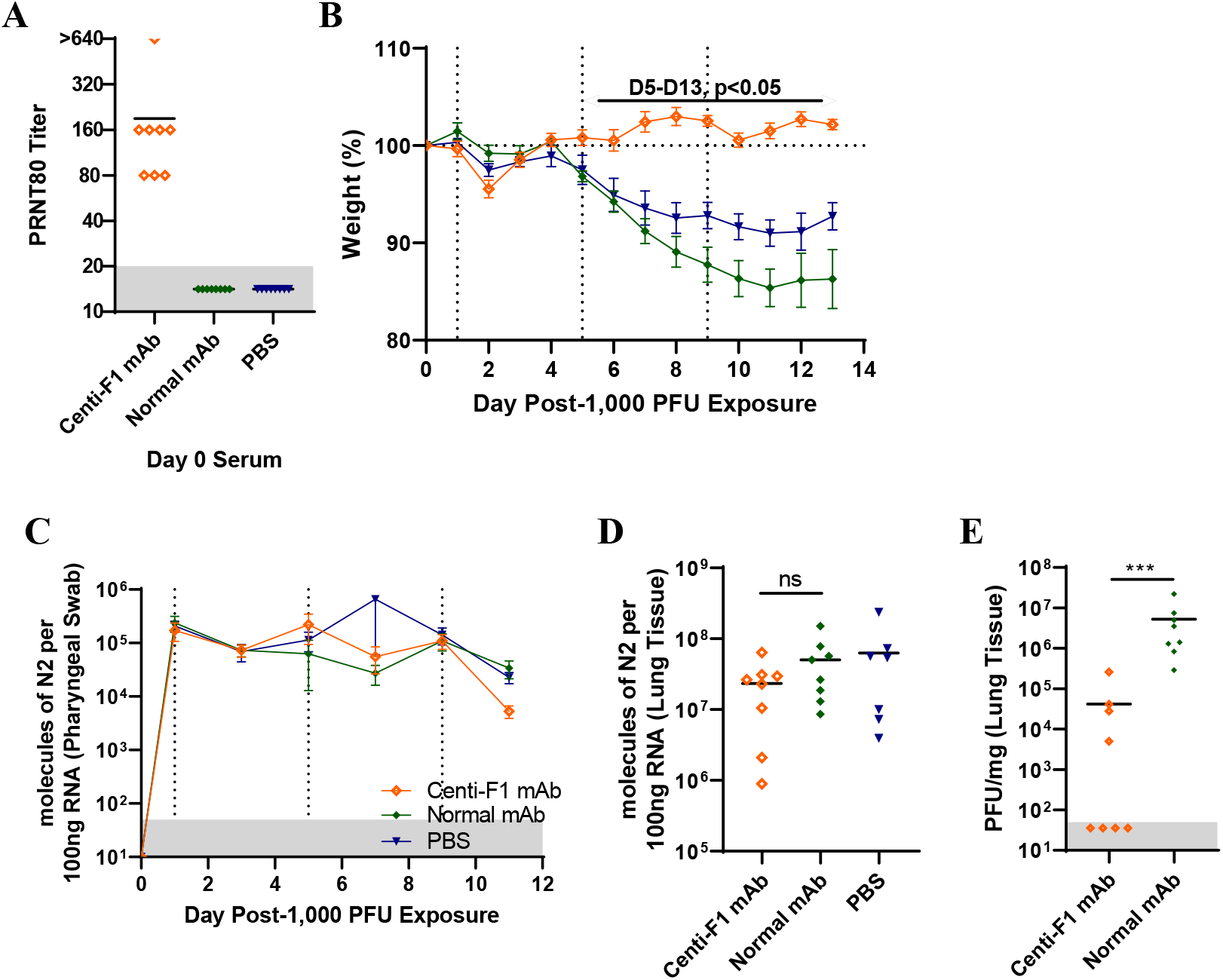
Passive transfer of anti-SARS-CoV-2 mAb Centi-F1 in immunosuppressed hamsters. Groups of 8 hamsters were immunosuppressed with CyP beginning −3 dpi (and indicated by the vertical lines in **B, C**) and passively transferred 30 mg/kg of Centi-F1 mAb, equivalent volume of normal mAb or PBS −1 dpi and exposed to 1,000 PFU SARS-CoV-2 on Day 0. **A)** Levels of circulating neutralizing antibody from Day 0 serum was assayed by PRNT. Disease progression was monitored by **B)** weight and **C)** pharyngeal swabs. Lung tissue collected 13 dpi was assayed for **D)** viral RNA and **E)** infectious virus.

## Discussion

It has been postulated that both innate immune hyperactivity and adaptive immune dysfunction result in increased SARS-CoV-2 dissemination and drives severe disease (*14*). In order to develop a small animal model of severe disease, evaluate the role of adaptive immunity, and replicate the COVID-19 lymphopenia observed in human disease, we treated hamsters with CyP. In this immunosuppressed model, we observe more drastic weight loss, high levels of virus in the lung, and viral persistence that did not begin to resolve until CyP treatment was stopped (**Fig. 1**). Histopathologic findings in the lungs of CyP-treated hamsters also demonstrate the prolonged pathology and ongoing repair through hyperplastic changes of the bronchial epithelium and type II pneumocytes that were not observed in the sampled *RAG2* KO hamsters. These data demonstrate the critical role B and T cells play in resolution of disease, and describe an animal model for severe COVID-19 disease that can be used for rigorous testing of medical countermeasures.

The recent development of immune function KO hamsters combined with Syrian hamster susceptibility to SARS-CoV-2 infection allows investigation of the role of a specific immune functions in COVID-19 pathogenesis and protection. Thus far, *STAT2* KO hamsters have been used to demonstrate that type I interferon plays a role in restricting viral dissemination and promoting lung pathology in SARS-CoV-2 infected animals (*15*). Herein we found, unexpectedly, that SARS-CoV-2 challenge resulted in a uniform lethality in the *RAG2* KO hamsters. It is interesting that onset of disease, as measured by weight loss, was more rapid (by 3 days) in the *RAG2* KO animals than immunocompetent, or even transiently immunosuppressed, hamsters receiving the same challenge dose. This indicates that the absence of functional B cells and/or T cells exacerbates pathogenesis at a very early stage (within a day or two) after exposure to virus. The importance of T cells in viral clearance has been noted for other coronaviruses, SARS-CoV (*16*) and MERS (*17*). In fact, depletion of CD3+ T cells correlates with severity and adverse outcomes (*18*).

There have been anecdotal accounts suggesting that immunity generated from an initial SARS-CoV-2 infection in humans may not be protective against a subsequent infection. Recent studies using nonhuman primates indicated prior infection results in protective immunity (*19*). Here, we show that the immune response from an initial SARS-CoV-2 infection, including a robust neutralizing antibody response, is protective against re-challenge. We detected reduced levels of viral genome in pharyngeal swabs and lungs, however, despite circulating neutralizing antibody available at the time of challenge, virus genome was still detected seven days after re-challenge. Viral genome in pharyngeal swabs on Day 5 following re-challenge and in lung tissue on Day 7 could indicate that asymptomatic reinfection is possible. Prolonged viral shedding has been observed in human COVID-19 cases, with detectable neutralizing antibody titers (*20*). Alternatively, the genome detected in the swabs could be input virus that is slowly being cleared from the respiratory system.

Currently, convalescent plasma is an option to treat severe cases of COVID-19 (*21*). Another antibody-based options include manufactured polyclonal or mAbs that represent avenues to prevent or treat SARS-CoV-2 infections as a standardized product. Several groups have reported *in vitro* characterization of anti-SARS-CoV-2 neutralizing mAbs and a number of antibodies have been shown to protect in small animal models, including Syrian hamsters (*22, 23*). Here, pre-treatment with mAb Centi-F1 resulted in weight maintenance through the experiment, trending lower levels of viral genome detected in pharyngeal swabs and lung tissues, and most importantly, reduced levels of infectious virus detected in the lung.

The lack of reduced viral RNA detected by RT-PCR in the passive transfer experiment of F01-mAb treated animals highlights a potential confounding factor in determining the efficacy of mAb treatment. It is possible that the Centi-F1 mAb was not able to clear genomic RNA but were able to prevent the shedding of infectious virus. In addition to determining infectious viral loads by plaque assay, RT-PCR to detect subgenomic RNA indicative of a productive infection would help differentiate the persistence of input virus from productive infection.

Our results expand on the earlier findings that the Syrian hamster model is a suitable small animal model of COVID-19 (*2*). Transient and reversible immunosuppression using CyP can be used to increase the severity and duration of the disease state. Advantages of the CyP model are that the wild type hamsters are readily available, and would produce a normal immune response to vaccination. One disadvantage is that disruption of lymphocytes could confound the evaluation of therapeutics that target components of the immune response, or vaccines that require rapid mobilization of the adaptive response. For evaluation of those types of medical countermeasures, the wild type hamster would be preferred. This is the first report that SARS-CoV-2 is lethal in *RAG2* KO hamsters. For practical purposes, uniformly lethal models allow fewer numbers of animals to detect significant levels of protection facilitating rapid screening of candidate therapeutics. *RAG2* KO hamsters would not be suitable for the use in active vaccine studies but could be used for antibody passive transfer studies. For example, passive transfer of immune serum from nonhuman primate or human vaccine studies could be an approach to investigate mechanisms of protection. Overall, our hamster SARS-CoV-2 models with severe diseases clinically relevant to those of COVID-19 patients provide a platform for evaluating candidate medical countermeasures to combat the pandemic.

## Acknowledgments

We would like to acknowledge the tireless efforts of the Comparative Medicine Division, Histology Lab and Aerobiological Sciences technicians at USAMRIID. We thank Joshua Moore, Jimmy Fiallos, Steven Stephens, Leslie Klosterman, Lynda Miller, Jua Liu, April Babka, Neil Davis, and Dave Dyer for assistance with veterinary care, histology and molecular assays, and hematology.

## Author contributions

R.L.B., J.W.G., and J.W.H. designed the study. R.K.K., X.Z., J.A.W., performed the pathology and imaging analyses. L.M.P. and J.M.S. performed the *in vitro* assays. R.L., Y.L., and Z.W. provided the *RAG2* KO animals. D.G., S.Y., and J.G. provided the Centi-F1 mAb. R.L.B. and J.W.H. wrote the paper with all the coauthors.

## Data and materials availability

All data is available in the main text or the supplementary materials.

## Supplementary Materials

Materials and Methods

Figures S1-S5

Tables S1-S4

