## Supplemental for "Disruption of Adaptive Immunity Enhances Disease in SARS-CoV-2 Infected Syrian Hamsters"

**This PDF file includes:**

Materials and Methods
Figs. S1 to S5
Tables S1 to S4

### Materials and Methods

**Ethics**. Animal research was conducted under an IACUC approved protocol at USAMRIID (USDA Registration Number 51-F-00211728 & OLAW Assurance Number A3473-01) in compliance with the Animal Welfare Act and other federal statutes and regulations relating to animals and experiments involving animals. The facility where this research was conducted is fully accredited by the Association for Assessment and Accreditation of Laboratory Animal Care, International and adheres to principles stated in the Guide for the Care and Use of Laboratory Animals, National Research Council, 2011.

**SARS-CoV-2 stock.** An aliquot of the second passage of SARS-CoV-2 USA-WA-1/2020 was received from the CDC and propagated in ATCC Vero 76 cells (99% confluent) in EMEM containing 1% GlutaMAX, 1% NEAA, and 10% heat-inactivated fetal bovine serum at an MOI of 0.01. Supernatant was collected from cultures exhibiting characteristic CPE and clarified by centrifugation (10,000 g x 10 minutes). Clarified virus was subjected to the following specifications: Identification by SARS-CoV-2 RT-PCR assay, Quantification by agarose-based plaque assay, free from contaminants by growth of chocolate agar plates, endotoxin testing using Endosafe® nexgen-PTS, and mycoplasma using MycoAlert test kit, and genomic sequencing.

**Animal Procedures.** Wild type (females only, aged 6-8 weeks) or RAG2 KO (females and males, aged 11-12 weeks) hamsters (*Mesocricetus auratus*) were anesthetized by inhalation of vaporized isoflurane using an IMPAC6 veterinary anesthesia machine for the following procedures: intranasal challenge of virus, CyP intraperitoneal injections, pharyngeal swabs, and non-terminal blood collection. Intranasal instillation of SARS-CoV-2 was administered in a volume of 50µl for challenge doses of 100, 1,000 and 10,000 PFU, and 100µl for the challenge dose of 100,000 PFU. CyP treatment (Baxter, pharmaceutical grade) consisted of an initial loading dose of 140mg/kg, followed by maintenance doses of 100mg/kg on the days indicated by each experiment. Pharyngeal swabs in 0.5ml of complete media were used for virus detection to monitor infection and disease course in hamsters. Vena cava blood collection was limited to 7% of total blood volume per week. Terminal blood collection was performed by cardiac injection at the time of euthanasia. All work involving animals was performed in an animal biosafety level 3 (ABSL-3) laboratory.

**Anti-SARS-CoV-2 mAb.** F01 mAb was administered to hamsters at a dose of 30mg/kg by the subcutaneous route. F01 mAb was a kind gift from Distributed Bio, Inc.

**Viral RNA assay.** Following 3 freeze/thaws of frozen swabs in media, 250µl of media was removed and added to 750µl of Trizol LS. Approximately 200mg of organ tissue was homogenized in 1.0ml of Trizol using M tubes on the gentleMACS dissociator system on the RNA setting. RNA was extracted from Trizol LS or Trizol per manufacturer’s protocol. A Nanodrop 8000 was used to determine RNA concentration, which was then raised to 100ng/µl in UltraPure distilled water. Samples were run in duplicate on a BioRad CFX thermal cycler using TaqPath 1-step RT-qPCR master mix according to the CDC’s recommended protocol of 25°C for 2 minutes, 50°C for 15 minutes, 95°C for 2 minutes, followed by 45 cycles of , 95°C for 3 seconds and 55°C for 30 seconds. The forward and reverse primer and probe sequences are: 2019-nCoV_N2-F, 5’-TTA CAA ACA TTG GCC GCA AA-3’, 2019-nCoV_N2-R, 5’-GCG

CGA CAT TCC GAA GAA-3’, and 2019-nCoV_N2-P, 5’-ACA ATT TCC CCC AGC GCT TCA G-3’. The limit of detection for this assay is 50 copies.

**PRNT.** An equal volume of complete media (EMEM containing 10% heat-inactivated FBS, 1% Pen/Strep, 0.1% Gentamycin, 0.2% Fungizone, cEMEM) containing SARS-CoV-2 was combined with 2-fold serial dilutions of cEMEM containing antibody and incubated at 37°C in a 5% $CO_2$ incubator for 1 hour (total volume 222µl). 180 µl per well of the combined virus/antibody mixture was then added to 6-well plates containing 3-day old, ATCC Vero 76 monolayers and allowed to adsorb for 1 hour in a 37°C, 5% $CO_2$ incubator. 3mL per well of agarose overlay (0.6% SeaKem ME agarose, EBME with HEPES, 10% heat-inactivated FBS, 100X NEAA, 1% Pen/Strep, 0.1% Gentamycin and 0.2% Fungizone) was then added and allowed to solidify at room temperature. The plates were placed in a 37°C, 5% $CO_2$ incubator for 2 days and then 2mL per well of agarose overlay containing 5% neutral red and 5% heat-inactivated FBS is added. After 1 additional day in a 37°C, 5% $CO_2$ incubator, plaques were visualized and counted on a light box. PRNT80 titers are the reciprocal of the highest dilution that results in an 80% reduction in the number of plaques relative to the number of plaques visualized in the cEMEM alone (no antibody) wells.

**Plaque Assay.** Approximately 200mg of lung tissue was homogenized in 1.0mL of cEMEM using a gentleMACS M tubes and a gentleMACS dissociator on the RNA setting. Tubes were centrifuged to pellet debris and supernatants collected. Ten-fold dilutions of the samples were adsorbed to Vero 76 monolayers (200µl of each dilution per well). Following a 1 hour adsorption in a 37°C, 5% $CO_2$ incubator, cells were overlaid and stained identically as described for PRNT. The limit of detection for this assay is 50 plaque forming units (PFU).

**Hematology.** Whole blood collected in EDTA tubes was analyzed on an HM5 hematology analyzer on the DOG2 setting.

**Preparation of tissues for histology.** Tissues were fixed in 10% neutral buffered formalin, trimmed, processed, embedded in paraffin, cut at 5 to 6µm, and stained with hematoxylin and eosin (H&E).

***In situ* hybridization.** To detect SARS-CoV-2 genomic RNA in FFPE tissues, *in situ* hybridization (ISH) was performed using the RNAscope 2.5 HD RED kit (Advanced Cell Diagnostics, Newark, CA, USA) as described previously (*1*). Briefly, forty ZZ ISH probes targeting SARS-CoV-2 genomic RNA fragment 21571-25392 (GenBank #LC528233.1) were designed and synthesized by Advanced Cell Diagnostics (#854841). Tissue sections were deparaffinized with xylene, underwent a series of ethanol washes and peroxidase blocking, and were then heated in kit-provided antigen retrieval buffer and digested by kit-provided proteinase. Sections were exposed to ISH target probe pairs and incubated at 40°C in a hybridization oven for 2 h. After rinsing, ISH signal was amplified using kit-provided Pre-amplifier and Amplifier conjugated to alkaline phosphatase and incubated with a Fast Red substrate solution for 10 min at room temperature. Sections were then stained with hematoxylin, air-dried, and cover slipped.

**Immunofluorescence.** Formalin-fixed paraffin embedded (FFPE) tissue sections were deparaffinized using xylene and a series of ethanol washes. After 0.1% Sudan black B (Sigma) treatment to eliminate the autofluorescence background, the sections were heated in Tris-EDTA buffer (10mM Tris Base, 1mM EDTA Solution, 0.05% Tween 20, pH 9.0) for 15 minutes to reverse formaldehyde crosslinks. After rinses with PBS (pH 7.4), the section were blocked with PBT (PBS +0.1% Tween-20) containing 5% normal goat serum overnight at 4°C. Then the sections were incubated with rabbit anti-SARS-CoV Spike (1:200, Sino Biological, 40150-T62-COV2) or mouse anti-SARS-CoV NP (1:200, Sino Biological, 40143-MM05) antibodies and mouse anti-pan-cytokeratin (1:100, Santa Cruz Biotechnology, sc-8018), mouse anti-CC10 (1:100, Santa Cruz Biotechnology, sc-365992), mouse anti-E-cadherin (1:100, Thermo Fisher, 33-4000antibodies for 2 hours at room temperature. After rinses with PBT, the sections were incubated with secondary goat anti-rabbit Alexa Fluor 488 (1:500, Thermo Fisher) and goat anti-mouse Alexa Fluor 568 (1:500, Thermo Fisher) antibodies, for 1 hour at room temperature. Sections were cover slipped using the Vectashield mounting medium with DAPI (Vector Laboratories). Images were captured on a Zeiss LSM 880 confocal system and processed using ImageJ software.

**Transmission electron microscopy.** Fresh hamster lung and trachea were harvested after euthanasia and submerged in 2.5% glutaraldehyde and 2% paraformaldehyde in 0.1M sodium phosphate buffer for 1-3 hours and then placed in 4% paraformaldehyde for 14 or 21 days for viral inactivation. Samples were submerged in microchem prior to removal from containment suites. Tissue was trimmed and then rinsed with 0.1M sodium cacodylate buffer before post-fixing with 1% osmium tetroxide in 0.1M sodium cacodylate. After osmium fixation, the samples were rinsed with 0.1M sodium cacodylate buffer, followed by a water wash then subjected to uranyl acetate *en bloc*. Samples were washed with water then dehydrated through a graded ethanol series including 3 exchanges with 100% ethanol. Samples were further dehydrated with equal volume of 100% ethanol and propylene oxide followed by two changes of propylene oxide. Samples were initially infiltrated with equal volumes of propylene oxide and resin (Embed-812; EMS, Hatfield, PA) then incubated overnight in propylene oxide and resin. Next day, the samples were infiltrated with 100% resin embedded and oriented in 100% resin and then allowed to polymerize for 48 hours at 60ºC. 1 micron thick sections were cut from each tissue block, a region of interest for thin sectioning was chosen and 80nm thin sections were cut and collected on 200 mesh copper grids. Two grids from each sample was further contrast stained with 2% uranyl acetate and Reynold's lead citrate. Samples were then imaged on the Jeol 1011 TEM at various magnifications.

**Statistical analyses.** Statistical analyses were completed using GraphPad Prism 8. Weight data was analyzed using a one-way ANOVA with multiple comparisons for experiments with ≥3 groups; unpaired t-tests were used to analyze weight data for experiments with 2 groups. Comparisons of lymphocyte levels and lung viral load was assessed using unpaired t-tests. Significance of survival data was assessed using log-rank tests. In all analyses, $P<0.05$ is considered statistically significant.

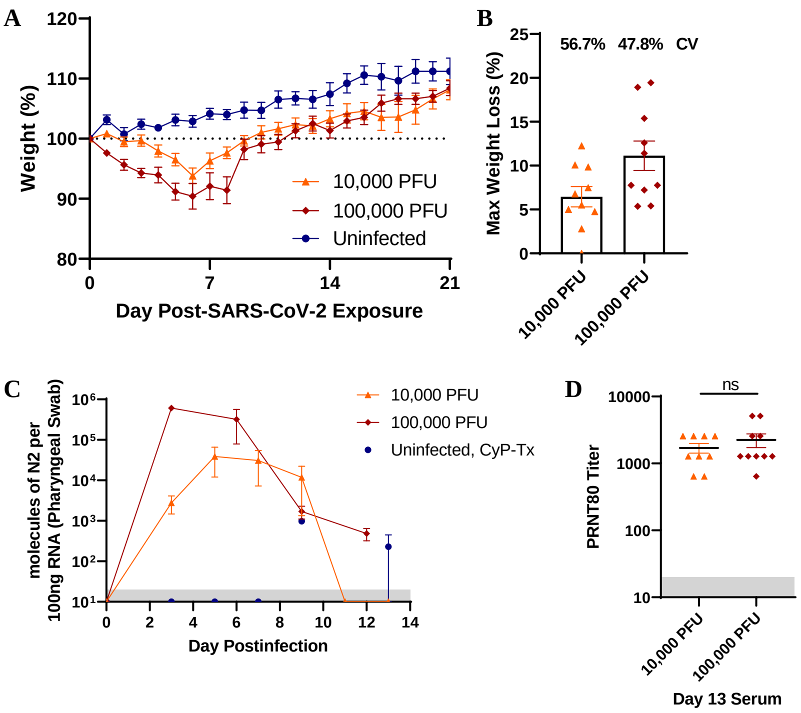


| **E** | Mean, SD, N, p-value compared to Uninfected, Significance | | |
|---|---|---|---|
| **Day** | **Uninfected** | **10,000 PFU** | **100,000 PFU** |
| D1 | 103.2%, 2.022, 6 | 100.8%, 1.076, 10, 0.3222, ns | 97.61%, 1.113, 10, <0.0001, **** |
| D2 | 100.7%, 2.672, 6 | 99.47%, 2.543, 10, 0.9774, ns | 95.66%, 2.907, 10, 0.0057, ** |
| D3 | 102.4%, 2.087, 6 | 99.67%, 3.030, 10, 0.3374, ns | 94.28%, 2.441, 10, 0.0002, *** |
| D4 | 101.8%, 1.148, 5 | 97.94%, 3.236, 10, 0.1120, ns | 93.96%, 4.161, 10, 0.0003, *** |
| D5 | 103.1%, 2.219, 5 | 96.51%, 3.353, 10, 0.0598, ns | 91.18%, 4.432, 10, 0.0002, *** |
| D6 | 102.9%, 2.136, 5 | 93.82%, 4.113, 10, 0.0198, * | 90.42%, 6.711, 10, 0.0009, *** |
| D7 | 104.2%, 2.071, 5 | 96.31%, 4.076, 10, 0.0509, ns | 92.10%, 7.073, 10, 0.0025, ** |
| D8 | 104.0%, 1.922, 5 | 97.67%, 3.062, 10, 0.0578, ns | 91.41%, 7.055, 10, 0.0005, *** |
| D9 | 104.7%, 2.981, 5 | 99.66%, 2.767, 10, 0.1326, ns | 98.21%, 5.408, 10, 0.0406, * |
| D10 | 104.7%, 3.050, 5 | 101.0%, 3.533, 10, 0.1880, ns | 99.09%, 4.570, 10, 0.0368, * |
| D11 | 106.5%, 3.188, 5 | 101.7%, 3.357, 10, 0.1056, ns | 99.43%, 4.007, 10, 0.0109, * |
| D12 | 106.7%, 2.457, 5 | 102.4%, 3.483, 10, 0.0971, ns | 101.3%, 3.692, 10, 0.0253, * |
| D13 | 106.6%, 3.283, 5 | 102.1%, 3.972, 10, 0.0971, ns | 102.5%, 3.930, 10, 0.1402, ns |

**Fig. S1. SARS-CoV-2-infected, immunocompetent hamsters have variable weight loss. A)** Groups of 10 each wild-type Syrian hamsters were infected with either 10,000 or 100,000 PFU SARS-CoV-2 by the intranasal route on Day 0 and weights monitored daily. **B)** The maximum weight loss for individual animals was plotted by group to determine the coefficient of variance. **C)** Pharyngeal swabs were used to monitor disease progression. **D)** Serum collected on Day 13 was evaluated for neutralizing antibody titer by PRNT. **E)** The mean, standard deviation, n are displayed for each group weight by day. A one-way ANOVA using multiple comparisons to Uninfected was calculated to determine p-value and significance.

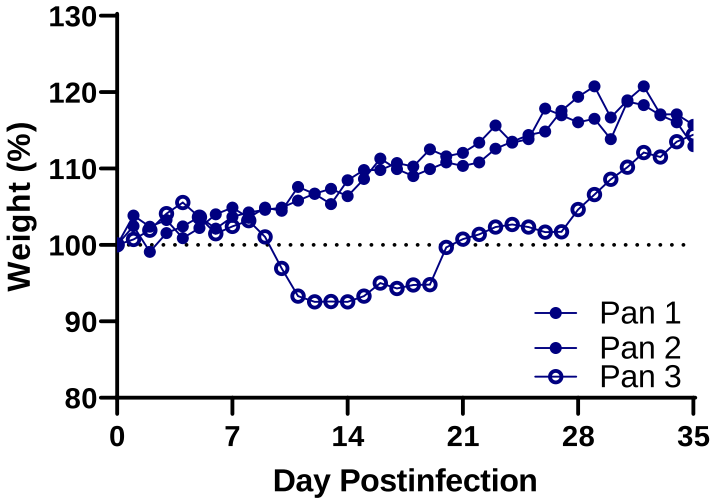


**Fig. S2. Transmission of SARS-CoV-2 to CyP-treated, mock-infected hamsters.** A single cage of 4 animals began losing weight on Day 9 relative to the other mock-infected hamsters from this group. These animals (Pan 3) were removed from weight data presented for uninfected animals in **Fig. 1.**

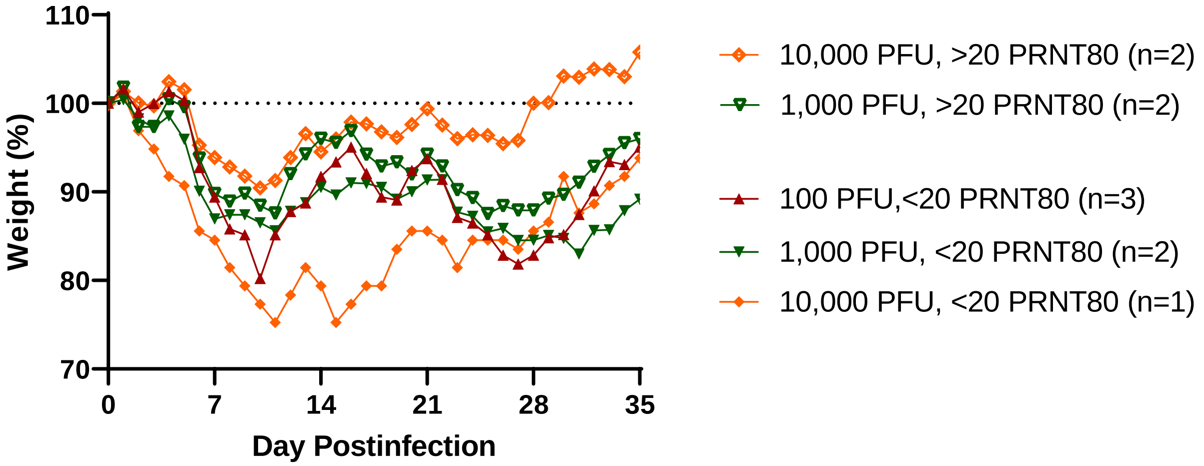


**Fig. S3. Weight kinetics of surviving hamsters groups according to positive/negative PRNT80 titers.** Weight data from surviving hamsters from the CyP-treated, SARS-CoV-2-infected groups from **Fig. 1** was groups according to challenge dose and <20 (closed symbols) or >20 (open symbols) PRNT80 titers.

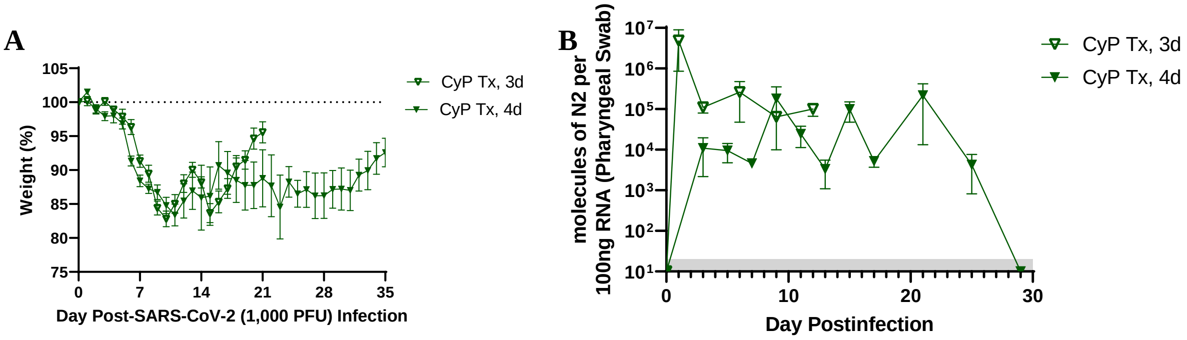


**Fig. S4. Increased frequency of CyP administration does not affect weight loss kinetics.** Groups of 10 hamsters each were administered CyP at either 3-day intervals (Days -3, 0, 3, 6, and 9 post-infection) or 4-day intervals (Days -3, 1, 5, 9, 13, 17, and 21 post-infection). **A)** Weights and **B)** pharyngeal swabs were used to monitor disease progression.

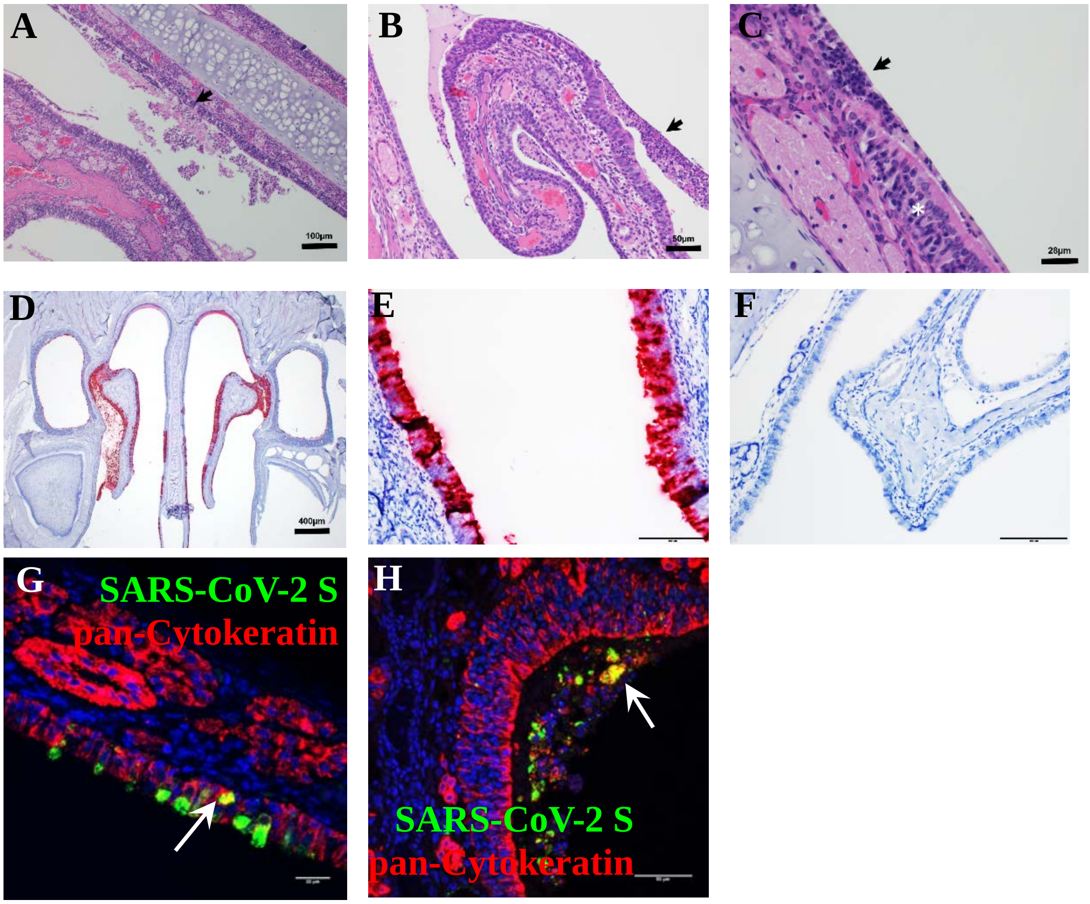


**Fig. S5. SARS-CoV-2 infects the epithelium of hamster nasal turbinates.** H&E sections of nasal turbinate tissue from SARS-CoV-2 infected, RAG2 KO hamsters show olfactory mucosal erosion and epithelial necrosis. **A-B)** Epithelial necrosis (arrow) overlaid by necrotic debris and degenerate inflammatory cells. The submucosa is expanded by inflammatory cells composed of heterophils and lymphocytes. **C)** Olfactory muscosal erosion and necrosis adjacent to intact less affected olfactory epithelium (asterisk). The mucosa is overlaid with a pseudomembrane composed of degenerate inflammatory cells, necrotic debris, and fibrin (arrow). Inflammatory cells extend into the submucosa. FFPE nasal turbinate tissue was stained for SARS-CoV-2 genomic RNA from SARS-CoV-2 exposed **D)** CyP-treated hamsters, **E)** RAG2 KO hamsters or **F)** untreated, uninfected hamsters. **G-H)** RAG2 KO hamster nasal turbinate tissue was stained for viral antigen (green) and epithelial cells (red) using antibodies against the SARS-CoV-2 spike protein and pan-cytokeratin. Infected epithelial cells are depicted by white arrows. Scale bars (**A**) = 100 microns; (**C**) = 28 microns; (**D**) 400 microns; (**E-H**) = 50 microns.

| | Mean, SD, N, p-value compared to Uninfected, CyP-Tx, Significance | | | |
|---|---|---|---|---|
| **Day** | **Uninfected, CyP-Tx** | **100 PFU, CyP-Tx** | **1,000 PFU, CyP-Tx** | **10,000 PFU, CyP-Tx** |
| D1 | 103.2%, 2.022, 6 | 101.9%, 1.660, 10, 0.1903, ns | 101.5%, 0.961, 10, 0.042, ** | 101.5%, 0.788, 10, 0.04, * |
| D2 | 100.7%, 2.672, 6 | 99.37%, 2.169, 10, 0.2909, ns | 99.0%, 1.914, 10, 0.152, ns | 99.33%, 3.340, 10, 0.392, ns |
| D3 | 102.4%, 2.087, 6 | 99.50%, 1.917, 10, 0.0128, * | 97.9%, 1.963, 9, 0.001, *** | 98.80%, 4.091, 10, 0.065, ns |
| D4 | 101.8%, 1.148, 5 | 100.1%, 2.583, 10, 0.1754, ns | 98.0%, 3.181, 9, 0.025, ** | 99.21%, 4.936, 10, 0.272, ns |
| D5 | 103.1%, 2.219, 5 | 98.80%, 2.533, 10, 0.007, *** | 96.9%, 2.424, 9, 0.0005, *** | 97.35%, 5.459, 10, 0.044, * |
| D6 | 102.9%, 2.136, 5 | 91.54%, 2.542, 10, <0.0001, **** | 91.34%, 2.026, 8, <0.0001, **** | 91.39%, 5.025, 10, 0.0003, ** |
| D7 | 104.2%, 2.071, 5 | 88.63%, 2.216, 10, <0.0001, **** | 88.42%, 2.403, 8, <0.0001, **** | 88.14%, 4.803, 10, <0.0001, **** |
| D8 | 104.0%, 1.922, 5 | 87.17%, 2.359, 10, <0.0001, **** | 87.24%, 1.925, 8, <0.0001, **** | 86.93%, 4.493, 10, <0.0001, **** |
| D9 | 104.7%, 2.981, 5 | 86.65%, 4.027, 10, <0.0001, **** | 86.74%, 3.017, 8, <0.0001, **** | 87.17%, 4.701, 10, <0.0001, **** |
| D10 | 104.7%, 3.050, 5 | 84.12%, 4.636, 10, <0.0001, **** | 84.79%, 3.400, 8, <0.0001, **** | 85.02%, 4.355, 10, <0.0001, **** |
| D11 | 106.5%, 3.188, 5 | 84.09%, 5.097, 10, <0.0001, **** | 83.39%, 4.524, 8, <0.0001, **** | 84.24%, 4.945, 10, <0.0001, **** |
| D12 | 106.7%, 2.457, 5 | 87.35%, 5.084, 10, <0.0001, **** | 85.45%, 7.193, 8, <0.0001, **** | 87.82%, 5.142, 10, <0.0001, **** |
| D13 | 106.6%, 3.283, 5 | 87.88%, 5.084, 10, <0.0001, **** | 86.96%, 7.811, 8, 0.0003, *** | 90.21%, 5.059, 10, <0.0001, **** |
| D14 | 107.4%, 3.847, 4 | 88.97%, 6.255, 7, 0.0005, *** | 85.95%, 11.688, 6, 0.008, ** | 89.13%, 6.130, 7, <0.0001, **** |
| D15 | 109.2%, 3.252, 4 | 89.40%, 6.453, 7, 0.0003, *** | 86.15%, 10.588, 6, 0.003, ** | 88.04%, 8.031, 7, 0.0005, *** |
| D16 | 110.6%, 3.097, 4 | 91.26%, 6.734, 7, 0.0005, *** | 90.69%, 7.862, 5, 0.002, ** | 89.45%, 8.517, 7, 0.0007, *** |
| D17 | 110.3%, 3.097, 4 | 88.26%, 6.520, 7, 0.0002, *** | 90.00%, 7.042, 5, 0.001, *** | 89.27%, 7.275, 7, 0.001, *** |
| D18 | 109.2%, 4.827, 4 | 86.74%, 7.001, 7, 0.0003, *** | 88.51%, 7.342, 5, 0.001, *** | 88.57%, 6.575, 7, 0.0006, *** |
| D19 | 111.2%, 3.969, 4 | 83.99%, 7.868, 7, 0.0001, *** | 87.77%, 8.142, 5, 0.001, *** | 89.37%, 5.430, 7, 0.0006, *** |
| D20 | 111.2%, 3.215, 4 | 84.16%, 11.83, 7, 0.002, *** | 87.75%, 7.680, 5, 0.0008, *** | 91.03%, 5.375, 7, 0.0008, *** |
| D21 | 111.2%, 4.447, 4 | 88.82%, 7.521, 7, 0.0007, *** | 88.77%, 9.378, 5, 0.003, ** | 91.82%, 6.373, 7, 0.0005, *** |
| D22 | 112.1%, 5.043, 4 | 89.15%, 6.496, 5, 0.0007, *** | 87.68%, 10.159, 5, 0.004, ** | 89.53%, 6.532, 7, 0.0002, *** |

| | | | | |
|---|---|---|---|---|
| D23 | 114.1%, 4.607, 4 | 84.36%, 8.208, 5, 0.0004, *** | 84.56%, 10.494, 5, 0.001, *** | 86.84%, 7.171, 7, 0.0008, *** |
| D24 | 113.4%, 3.071, 4 | 83.15%, 8.274, 5, 0.0002, *** | 88.28%, 4.533, 4, <0.00001, **** | 87.65%, 7.018, 7, 0.0001, *** |
| D25 | 114.1%, 4.754, 4 | 81,32%, 8.248, 5, 0.0002, *** | 86.51%, 3.940, 4, 0.0003, *** | 87.07%, 7.545, 7, 0.0001, *** |
| D26 | 116.4%, 6.195, 4 | 79.52%, 7.898, 5, <0.0001, **** | 87.14%, 5.258, 4, 0.0003, *** | 85.54%, 8.293, 7, 0.0001, *** |
| D27 | 117.3%, 3.710, 4 | 78.10%, 8.479, 5, <0.0001, **** | 86.22%, 6.707, 4, 0.0001, *** | 83.98%, 9.910, 7, 0.0001, *** |
| D28 | 117.8%, 3.279, 4 | 81.01%, 7.948, 4, 0.0001, *** | 86.24%, 6.687, 4, 0.0001, *** | 87.82%, 12.515, 5, 0.002, ** |
| D29 | 118.6%, 3.561, 4 | 82.25%, 8.736, 4, 0.0002, *** | 87.17%, 5.588, 4, 0.0001, *** | 87.31%, 13.096, 5, 0.003, ** |
| D30 | 115.3%, 3.346, 4 | 85.17%, 7.447, 3, 0.0007, *** | 87.23%, 6.245, 4, 0.0002, *** | 92.45%, 14,733, 4, 0.024, * |
| D31 | 118.9%, 2.373, 4 | 87.47%, 8.059, 3, 0.0006, *** | 87.01%, 5.971, 4, 0.0001, *** | 90.62%, 16.348, 3, 0.014, * |
| D32 | 119.5%, 2.108, 4 | 90.11%, 6.927, 3, 0.0004, *** | 89.26%, 4.731, 4, <0.00001, **** | 98.81%, 9.253, 3, 0.007, ** |
| D33 | 117.0%, 4.467, 4 | 93.41%, 4.676, 3, 0.001, *** | 89.26%, 5.634, 4, 0.0002, *** | 99.45%, 8.468, 3, 0.015, * |
| D34 | 116.6%, 3.508, 4 | 93.07%, 6.022, 3, 0.001, *** | 91.71%, 4.660, 4, 0.0001, *** | 99.27%, 6.828, 3, 0.007, ** |
| D35 | 114.3%, 4.909, 4 | 95.06%, 6.934, 3, 0.007, *** | 92.58%, 4.227, 4, 0.0005, *** | 101.79%, 7.153, 3, 0.039, * |

**Table S1. SARS-CoV-2/CyP-Tx Statistics.** For the weight data presented in **Fig. 1,** the mean, standard deviation, n are displayed for each group by day. A one-way ANOVA using multiple comparisons to Uninfected CyP-Tx was calculated to determine p-value and significance.

| | Mean, SD, N, p-value compared to RAG2 KO, Significance | | |
|---|---|---|---|
| **Day** | **RAG2 KO** | **CyP-Tx** | **CyP-Tx, Uninfected** |
| D1 | 98.44%, 1.007, 7 | 101.5%, 0.7885, 10, 0.0046, ** | 103.2%, 2.022, 6, 0.0007, *** |
| D2 | 97.53%, 1.468, 7 | 99.33%, 3.340, 10, 0.4051, ns | 100.7%, 2.672, 6, 0.0829, ns |
| D3 | 92.29%, 2.903, 6 | 98.80%, 4.091, 10, 0.0199, * | 102,4%, 2.087, 6, 0.0017, ** |
| D4 | 90.17%, 3.181, 6 | 99.21%, 4.936, 10, 0.0209, * | 101.8%, 1.148, 5, 0.0074, ** |
| D5 | 87.04%, 3.887, 6 | 97.35%, 5.459, 10, 0.0316, * | 103.1%, 2.219, 5, 0.0009, ** |
| D6 | 84.25%, 4.045, 6 | 91.39%, 5.025, 10, 0.1610, ns | 102.9%, 2.136, 5, 0.0064, *** |
| D7 | 83.59%, 4.775, 3 | 88.14%, 4.803, 10, >0.9999, ns | 104.2%, 2.071, 5, 0.0124, * |
| D8 | 82.90%, 3.012, 2 | 86.93%, 4.493, 10, 0.7148, ns | 104.0%, 1.922, 5, 0.0130, * |

**Table S2. SARS-CoV-2/RAG2 KO Statistics.** For the weight data presented in **Fig. 2,** the mean, standard deviation, n are displayed for each group by day. A one-way ANOVA using multiple comparisons to RAG2 KO was calculated to determine p-value and significance. Analysis was not conducted beyond D8 since only a single hamster remained in the RAG2 KO group.

Weights

| | Mean, SD, N, p-value, Significance | |
|---|---|---|
| **Day** | **Rechallenge** | **Naïve** |
| D44 | 98.58%, 1.293, 9, 0.5040, ns | 98.09%, 1.566, 7 |
| D45 | 100.1%, 1.347, 9, <0.0001, **** | 93.98%, 2.428, 7 |
| D46 | 101.5%, 2.556, 9, 0.0001, *** | 92.83%, 4.029, 7 |
| D47 | 101.6%, 2.311, 9, <0.0001, **** | 91.86%, 4.286, 7 |
| D48 | 102.2%, 2.817, 9, <0.0001, **** | 89.48%, 4.927, 7 |
| D49 | 103.0%, 2.466, 9, <0.0001, **** | 88.25%, 6.915, 7 |
| D50 | 103.6%, 3.023, 9, 0.0001, *** | 88.12%, 9.608, 7 |

Pharyngeal Swabs

| | Mean, SD, N, p-value, Significance | |
|---|---|---|
| **Day** | **Rechallenge** | **Naïve** |
| D44 | 247234, 241258, 7, 0.0023, ** | 2382049, 1558120, 6 |
| D46 | 16902, 33769, 6, 0.0303, * | 527461, 807434, 5 |
| D48 | 4026, 7303, 5, 0.0498, * | 20828, 27348, 6 |

**Table S3. SARS-CoV-2 Rechallenge Statistics.** For the weight and pharyngeal swab data presented in **Fig. 5,** the mean, standard deviation, n are displayed for each group by day. Rechallenge group data was normalized to 100% on Day 43 (time of challenge) for statistical comparison to naïve group. An unpaired t test was calculated to determine p-value and significance.

CyP-Tx Hamster Weights

| | Mean, SD, N, p-value compared to Normal mAb, Significance | | |
|---|---|---|---|
| **Day** | **Centi-F1** | **PBS** | **Normal mAb** |
| D1 | 99.67%, 2.308, 8, 0.0668, ns | 100.3%, 1.232, 8, 0.1376, ns | 101.5%, 2.463, 8 |
| D2 | 95.55%, 2.579, 8, 0.0085, ** | 97.49%, 1.861, 8, 0.6174, ns | 99.21%, 2.314, 8 |
| D3 | 98.54%, 2.205, 8, >0.9999, ns | 98.33%, 1.456, 8, >0.9999, ns | 99.13%, 2.463, 8 |
| D4 | 100.6%, 1.936, 8, >0.9999, ns | 98.93%, 3.105, 8, 0.4432, ns | 100.4%, 0.910, 8 |
| D5 | 100.8%, 2.325, 8, 0.0369, * | 97.51%, 4.236, 8, >0.9999, ns | 96.84%, 1.556, 8 |
| D6 | 100.5%, 3.132, 8, 0.0075, ** | 94.93%, 4.817, 8, >0.9999, ns | 94.25%, 3.070, 8 |
| D7 | 102.4%, 3.010, 8, 0.0013, ** | 93.59%, 4.599, 8, >0.9999, ns | 91.23%, 3.629, 8 |
| D8 | 103.0%, 2.665, 8, 0.0002, *** | 92.56%, 4.160, 8, >0.9999, ns | 89.10%, 4.496, 8 |
| D9 | 102.5%, 1.568, 8, <0.0001, **** | 92.81%, 3.551, 7, 0.6636, ns | 87.76%, 5.091, 8 |
| D10 | 100.6%, 2.085, 8, <0.0001, **** | 91.67%, 3.501, 7, 0.4992, ns | 86.35%, 5.230, 8 |
| D11 | 101.5%, 2.267, 8, <0.0001, **** | 91.01%, 3.580, 7, 0.6731, ns | 85.39%, 5.466, 8 |
| D12 | 102.7%, 2.181, 8, 0.0002, *** | 91.17%, 5.031, 7, >0.9999, ns | 86.16%, 7.826, 8 |
| D13 | 102.2%, 1.481, 8, 0.0002, *** | 92.74%, 3.694, 7, 0.7788, ns | 86.28%, 8.560, 8 |

CyP-Tx Hamster Swabs

| | Mean, SD, N, p-value compared to Normal mAb, Significance | | |
|---|---|---|---|
| **Day** | **Centi-F1** | **PBS** | **Normal mAb** |
| D1 | 172262, 129402, 4, >0.9999, ns | 208023, 94608, 4, >0.9999, ns | 237359, 157029, 4 |
| D3 | 73325, 38121, 4, >0.9999, ns | 68934, 49361, 4, >0.9999, ns | 73115, 35954, 4 |
| D5 | 219640, 250951, 4, 0.6536, ns | 113553, 90488, 4, 0.3069, ns | 62785, 99676, 4 |
| D7 | 55551, 57924, 4, 0.8655, ns | 660499, 1039780, 3, 0.1143, ns | 27156, 21819, 4 |
| D9 | 109814, 67872, 4, >0.9999, ns | 141341, 99141, 4, >0.9999, ns | 109846, 77393, 4 |
| D11 | 5285, 2760, 4, 0.1038, ns | 22677, 9296, 3, >0.9999, ns | 33881, 25010, 4 |

**Table S4. SARS-CoV-2 Passive Transfer Statistics.** For the weight and pharyngeal swab data presented in **Fig. 6,** the mean, standard deviation, n are displayed for each group by day. An unpaired t test was calculated to determine p-value and significance.
